# Predicting Dengue Fever in Brazilian Cities

**DOI:** 10.1101/2021.02.17.430949

**Authors:** Kirstin Roster, Colm Connaughton, Francisco A. Rodrigues

**Affiliations:** Institute of Mathematics and Computer Science, University of São Paulo, São Carlos, Brazil; Mathematics Institute, University of Warwick, Coventry, CV4 7AL, United Kingdom & London Mathematical Laboratory, 8 Margravine Gardens, London, W6 8RH, United Kingdom

## Abstract

Dengue Fever is an increasingly serious public health concern both in Brazil and globally. In the absence of a universal vaccine or specific treatments, prevention relies on vector control and disease surveillance. Accurate and early forecasts can help reduce the spread of the disease. In this study, we develop a model to predict the number of Dengue Fever cases in Brazilian cities one month ahead. We compare different machine learning approaches as well as different sets of input features based on epidemiological and meteorological data. We find that different models work best in different cities, and a random forests model trained on data of historical Dengue cases performs best overall. It produces lower aggregate errors than a seasonal naïve baseline model, Gradient Boosting Regression, feed-forward Neural Networks, and Support Vector Regression. Predictions on an unseen test set are on average within 11.5 cases for the median city. Mean absolute errors on the hold-out test set are reduced to 10.8 for the median city when selecting the optimal combination of algorithm and input features for each city individually.

## I. INTRODUCTION

Dengue Fever is a serious public health concern, burdening individuals’ lives and national economies. It is a mosquito-borne infectious disease that affects 100 to 400 million people each year [1]. Half of the world’s population and 129 countries are at risk of infection [2].

Brazil is home to about half of all reported cases of Dengue infection in the Americas. In the three decades after 1986, when official national reporting began, over eleven million suspected cases and over five thousand confirmed deaths were reported. Both infection and fatality rates increased over this period. The Southeastern and Northeastern regions are particularly affected, but nearly all states counted Dengue-related deaths. Several outbreaks have occurred, primarily due to re-introduction of one of the four Dengue serotypes. In 2007, the serotype DENV-2 re-emerged, causing an epidemic that disproportionately affected children, who accounted for over half of the epidemic’s deaths [3]. Besides human cost, Dengue has a significant impact on the Brazilian economy. During the 2013 outbreak alone, the economic burden was estimated at three hundred million USD [4].

The disease burden is expected to rise in light of global changes in climate [5], increasing deforestation, and disruption of natural ecological systems [6–9]. In Brazil specifically, deforestation has been linked to a recent outbreak of the Zika virus [10], which is transmitted by the same mosquito species as Dengue Fever. Another important risk factor in Brazil is the high number of hydro-electric dams, which also alter local ecological and social systems [11] and whose construction is correlated with reemergence of Malaria and other diseases [7, 12].

There is no vaccine against Dengue and few treatment options exist. Prevention relies on vector control, which underscores the importance of disease surveillance [13]. To best inform public health decisions such as resource allocation, disease forecasts need to be available early and at a granular geo-spatial resolution, while maintaining a high level of accuracy. Yet Dengue Fever is difficult to predict. Infection with one of the four Dengue serotypes provides lifetime immunity to that serotype as well as temporary cross-immunity to other serotypes, resulting in irregular periodicity of outbreaks. Dengue incidence is also influenced by a wide range of factors, including climate conditions, human mobility [14], and land use [15]. The relationship between Dengue and climate in particular has been extensively studied, and associations have been found with rainfall [16, 17], climate change [5, 18], temperature [16, 17, 19, 20], extreme weather events such as El Niño and La Niña [16, 20], humidity [19], atmospheric pressure [20], and sea surface temperatures [17, 21]. These factors may have nonlinear, context-specific, and time-variant effects on disease incidence, which poses another challenge to disease modeling.

The existing literature on Dengue covers a range of forecasting approaches, including both theoretical and data-centric methods. The classical epidemiological approach is compartmental modeling, which assesses the evolution of the number of infectious individuals and other compartments within a population (for example [22, 23]). Agent-based models simulate the actions of individuals, and can provide geo-spatial estimates of the spread of disease. They are helpful in assessing the potential impacts of different public policies that may alter individual behaviors or environmental conditions, such as the release of sterile mosquitoes (for example [24, 25]). Yet these theoretical models tend to require significant knowledge of the disease, hosts, and transmission processes for parameter estimation. Statistical time series forecasting tools, such as the Seasonal Autoregressive Integrated Moving Averages (SARIMA) model, employ a more data-centric approach and leverage the highly auto-correlated nature of Dengue Fever to predict future incidence (for example [26, 27]).

Machine learning models take advantage of the increasing measurement capacity and public availability of epidemiological and other data sources, and often incorporate novel big data streams, such as social media activity, mobile phone data, or search engine queries. Their advantage relative to other forecasting approaches is their ability to estimate parameters directly from the data. Neural networks in particular have demonstrated a powerful forecasting ability for diseases such as Malaria [28], Influenza [29], and Covid-19 [30].

Machine learning models have also been applied to forecast Dengue Fever in different contexts, often incorporating climate data. Baquero et al (2018) [31], for example, compare the performance of machine learning and statistical models to forecast the number of Dengue cases in the city of São Paulo in Brazil. In [32], Dengue risk is assessed even more granularly in Rio de Janeiro, another Brazilian city. The authors use Convolutional Neural Networks (CNN) based on aerial and street view images to predict Dengue risk at the neighborhood level, which is correlated with the real incidence rate. The literature shows that the most effective machine learning model for Dengue prediction varies across contexts, such as data inputs or location. In [33], the authors use epidemiological, climate and Baidu search data to forecast the number of Dengue cases in Guangdong province in China. They compare several different algorithms, and achieve the best results with a Support Vector Regression (SVR) model. Xu et al. (2020) [34] developed a recurrent neural network model with a Long Short-Term Memory (RNN-LSTM) layer to predict Dengue in 20 Chinese cities. Their RNN-LSTM model outperforms a SVR model. They also show the utility of a recent development in machine learning research called transfer learning, where the model is trained on data from a city with many Dengue cases and then used to predict cases in a different city with fewer cases.

In this study, we assess the potential of different machine learning models in predicting Dengue Fever one month ahead in over two hundred Brazilian cities. We compare different algorithms, including decision tree ensemble approaches, neural networks, support vector regression, and a seasonal naïve baseline. We also compare different sets of meteorological and epidemiological features to better understand the predictive contribution of different variables. The best model for each city was selected using expanding time series cross-validation and tested on a hold-out test set. Best performance was achieved by the random forests algorithm. While climate variables improved predictions in some cities, the most important predictors across all cities were historical Dengue case counts. Our model outperforms existing comparable approaches by achieving a lower forecast error for São Paulo than [31].

## II. MATERIALS AND METHODS

### A. Data

We use official government sources for epidemiological and meteorological variables. Monthly Dengue cases are reported for each Brazilian municipality in the *Sistema de Informação de Agravos de Notificação* (SINAN) data system [35] for the years spanning 2007-2017. The *Instituto Nacional de Meteorologia* (INMET) provides meteorological data for weather stations across Brazil [36]. Daily data is collected from 1/1/2005 until 31/12/2017 in accordance with the availability of epidemiological data from SINAN. Daily climate records are aggregated to monthly time series using both the mean and standard deviation to account for the overall quantity as well as the variability in climate. The included variables are (i) rainfall, (ii) maximum temperature, (iii) minimum temperature, (iv) relative median temperature, (v) insolation, which is the amount of solar energy reaching the earth, (vi) rate of evaporation (Piche), (vii) median relative humidity, and (viii) median wind speed.

We merge the two data sources using the coordinates of the weather stations and the geographic boundaries of the municipalities. After removal of missing data, the combined dataset covers 234 municipalities. Data is split into a training set (9 years, Jan 2007 - Dec 2015) and a hold-out test set (2 years, Jan 2016-Dec 2017). The data is normalized to have a mean of zero and a standard deviation of one. The normalization is performed using the training data only. During cross-validation, the mean and standard deviation are computed separately on each training fold.

### B. Feature Selection

We compare four different sets of input features, two of which are selected naïvely, and two of which are selected using relationship metrics.

- The first set contains only the past eleven lags of Dengue cases.
- The second set of features also includes eleven months of all climate variables in addition to eleven months of Dengue cases. As we use eight different climate variables, aggregated to monthly time series using two statistics, and eleven lags of each, we have a total of 187 epidemiological and meteorological features.
- We compare two feature selection approaches to reduce the relatively high dimensionality of inputs. The third set of features is determined using a causal approach to feature selection based on the PCMCI causal discovery algorithm. A total of seven features are selected at significance level *α* = 0.05.
- The fourth set of features is made up of the variables that are most strongly correlated with the number of Dengue cases. The number of features is fixed to seven, so as to allow for direct comparison with the PCMCI approach.

PCMCI [37] is a two-stage causal discovery algorithm for high-dimensional time series data. In the first stage, using a modified version of the *PC* algorithm [38] called *PC*_1_, iterative conditional independence tests are performed to identify relevant conditions for all variables. For each variable 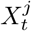 with the set of parents 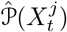, the variable 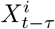 is removed from 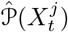 if 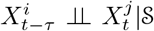, where 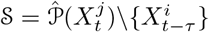, cannot be rejected at a given significance level *α*_*PC*_. Different kinds of independence tests can be performed at this stage, including the partial correlation test (ParCorr), which was implemented in this study. The second stage of PCMCI filters out false positives from each variable’s set of parents using the momentary conditional independence (*MCI*) test (formula 1):

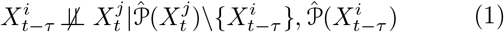

*MCI* assesses whether a variable 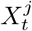 is independent of any of the parents, 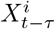, identified during the *PC*_1_ stage, conditional on both the remaining set of its parents, 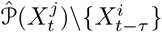 and the (time-shifted) parents of 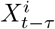, i.e. 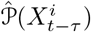.

### C. Prediction

We compare the following forecasting algorithms. As a baseline, we implement a seasonal naïve model (s-naïve) [39], which predicts the number of cases *y* in a given month *t* to be equal to the number of cases that occurred in the same month of the previous year: *ŷ*_*t*_ = *y*_*t*−12_. We compare the baseline to the following machine learning algorithms.

Random Forests (RF) [40] uses an ensemble of decision trees to make predictions. Each tree is grown by selecting a subset of observations in the training set with replacement (bootstrap aggregating, i.e. bagging) and then determining their best split based on a random subset of features using the Mean Squared Error (MSE) as splitting criterion.

Another tree-based ensemble approach is Gradient Boosting Regression (GBR) [41]. As with RF, final predictions are determined by the combined results of several decision trees. Unlike RF, GBR builds trees one at a time, applying boosting at each iteration. Boosting is the process of giving a higher weight to examples that are difficult to predict, thus incentivizing the model to improve its forecasts for the examples it predicted incorrectly previously.

Support Vector Regression (SVR) [42, 43] is a statistical learning technique that first transforms the input space using a kernel function, and then fits a linear function on the data. We use the radial basis function as the kernel. SVR is *ϵ*-insensitive, meaning errors of absolute magnitude up to *ϵ* are ignored, but errors that fall outside this range are minimized, while at the same time maintaining flatness of the fitted function (equation 2):

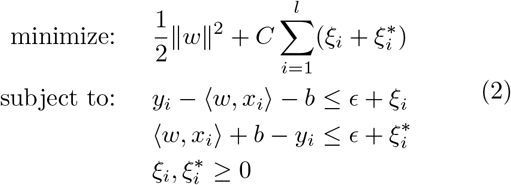

where *w* are weights, *C* is a constant to determine the flatness, (*x*_*i*_, *y*_*i*_) are the pairs of training feature vectors and corresponding targets, 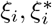 measure the cost of errors that is greater than |*ϵ*|, and *b* is the bias constant [44, 45].

Finally, we implement a multi-layer perceptron (MLP) neural network, comparing different architectures of up to three layers with up to 128 hidden units. Feed-forward neural networks consist of two stages. First, in the forward propagation step (equation 3), each neuron transforms inputs *x*_*j*_ to activations *a*_*k*_ using a set of weights *w* and biases *b* as well as a nonlinear activation function *g*:

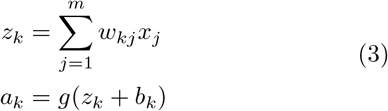

where *w*_*kj*_ is a matrix of weights, *b*_*k*_ is the bias vector, and *g* is the activation function.

The activations are passed as inputs to the neurons in the next layer of the network until they reach the final layer. The output of the last layer is a prediction *ŷ*, which is compared to the true training labels using a loss function.

During the backpropagation stage, the network parameters are adjusted to minimize the prediction errors. Learning consists of iteratively updating the weights and biases in the network using gradient descent as follows (equation 4):

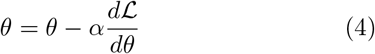

where *θ* is the parameter to be updated, ℒ is the loss, and *α* is the learning rate [46]. We use a Rectified Linear Unit (ReLU) activation function, Adam optimization function, 500 epochs, learning rate of 0.001, MSE loss function, and minibatch size of 200 observations. To avoid overfitting our predictions to the training examples, we implement L2 regularization with a parameter of 0.2.

We tested a range of hyperparameters for each of the machine learning algorithms, which is presented in table I.

**TABLE I.** Hyperparameters

| Algorithm | Hyperparameter | Values |
| --- | --- | --- |
| RF | number of trees: | [50, 100, 150, 200] |
|  | maximum tree depth: | [2,3,6, None] |
| GBR | number of trees: | [50, 100, 150, 200] |
|  | maximum tree depth: | [2,3,6, None] |
| MLP | number of layers: | [1, 2, 3] |
|  | number of units per layer: | [32, 64, 128] |
| SVR | epsilon: | [0.01, 0.05, 0.1, 0.2] |

### D. Model Evaluation

We use the Root Mean Square Error (RMSE) (equation 5) and the Mean Absolute Error (MAE) (equation 6) for model evaluation and selection:

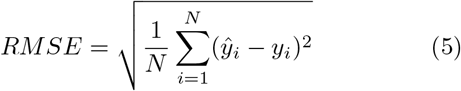

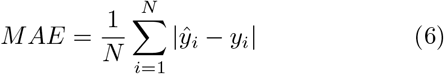

where *N* is the total number of observations, *ŷ* are the predicted values, and *y* are the actually observed number of cases.

Models are selected using expanding time series crossvalidation (CV). As we are working with sequence data, we cannot implement traditional CV, which would involve randomly shuffling the data to split it into training and validation sets. Instead, we maintain the chronological order of the data, by shifting our validation fold for each iteration of CV. We begin with one year of training data (year 2007) and the following year of validation data (year 2008). We then shift the validation fold to the next year (2009) and use all previous observations for training (years 2007-08). This shifting process is repeated until the full training data is covered, resulting in eight separate validation folds. The validation error of each city is computed as an average across the eight folds.

## III. RESULTS

At the validation stage, we use expanding time series CV to select the optimal hyperparameters for each model, and compare the relative performance of each combination of algorithm and feature set. The selected hyperparameter values vary across the sets of input features and are presented in table II.

**TABLE II.**
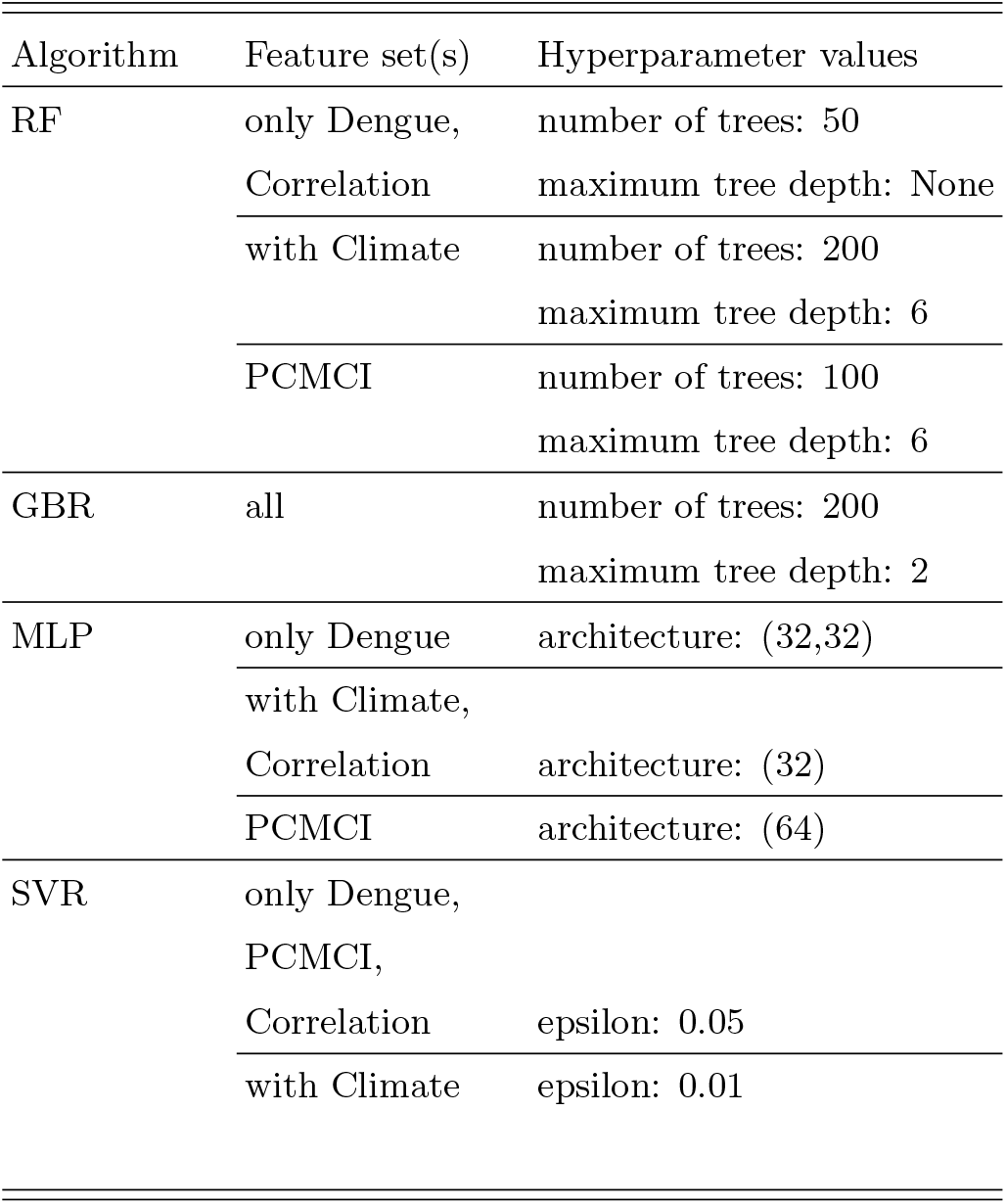
Optimal hyperparameters selected using CV

The Random Forests (RF) algorithm performed best on the validation folds, especially with only Dengue inputs. This combination of algorithm and feature set had the lowest errors across all cities according to the mean MAE and RMSE. For the median city, this resulted in a RMSE of 15.5 and predictions on average within 8.6 cases (table III). PCMCI feature selection gave the lowest median MAE in combination with the RF algorithm. The lowest median RMSE was achieved by the GBR model trained on only Dengue inputs. When selecting a single model for each city, RF was chosen most frequently (figure 1). This result holds regardless of the error metric chosen as the minimizing criterion.

**TABLE III.** Validation errors

| Algorithm | Feature Set | MAE |  | RMSE |  |
| --- | --- | --- | --- | --- | --- |
|  |  | mean | median | mean | median |
| GBM | Correlation | 39.0 | 8.8 | 68.1 | 13.6 |
|  | PCMCi | 38.7 | 8.9 | 67.6 | 13.9 |
|  | only Dengue | 38.3 | 8.9 | 67.3 | **13.5** |
|  | with Climate | 41.4 | 9.7 | 72.5 | 15.6 |
| MLP | Correlation | 44.0 | 12.1 | 74.6 | 17.7 |
|  | PCMCi | 41.7 | 12.4 | 71.2 | 18.7 |
|  | only Dengue | 48.2 | 13.0 | 78.6 | 18.3 |
|  | with Climate | 53.8 | 22.7 | 89.9 | 32.0 |
| RF | Correlation | 37.4 | 8.9 | 67.4 | 15.4 |
|  | PCMCi | 38.8 | **8.5** | 67.9 | 14.2 |
|  | only Dengue | **37.2** | 8.6 | **67.3** | 15.5 |
|  | with Climate | 42.6 | 9.6 | 73.5 | 15.6 |
| SVR | Correlation | 54.6 | 15.1 | 90.5 | 18.0 |
|  | PCMCi | 54.6 | 15.3 | 90.4 | 18.8 |
|  | only Dengue | 55.3 | 14.9 | 90.5 | 17.7 |
|  | with Climate | 53.0 | 14.8 | 91.1 | 19.9 |
| s-naïve |  | 80.8 | 14.1 | 206.4 | 31.2 |

Lowest errors are highlighted.

**FIG. 1.**
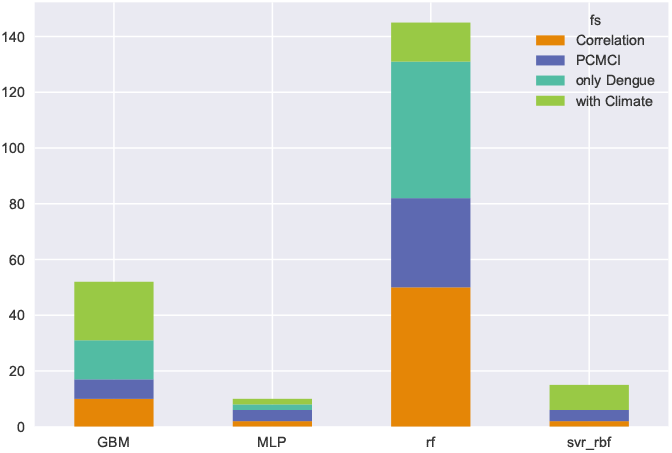
Distribution of algorithm and feature set among best model per city (lowest MAE)

The models selected on the validation sets are trained on the full training set and evaluated on the hold-out test set. The best overall model, the RF algorithm trained only on Dengue inputs, predicts Dengue Fever with a MAE of 11.5 for the median city on unseen data (table IV). Performance is improved slightly when selecting different combinations of algorithms and input features for each city individually, resulting in a MAE of 10.8 cases for the median city, as well as lower mean and median RMSE. Both approaches significantly outperform the naïve baseline forecast.

**TABLE IV.** Test set errors

| Model | MAE |  | RMSE |  |
| --- | --- | --- | --- | --- |
|  | mean | median | mean | median |
| RF (only Dengue) | **59.4** | 11.5 | 133.7 | 23.8 |
| City-specific | 62.2 | **10.8** | **129.9** | **18.8** |
| s-naïve | 174.4 | 17.3 | 324.7 | 30.4 |

Lowest errors are highlighted.

Qualitatively, the model is able to capture different kinds of time series, as illustrated in three sample cities in figure 2. São Paulo experiences annual Dengue outbreaks, though the total number of cases differs across years. Some years (e.g. 2014-16) have much larger peaks than the other years. Sorocaba is a special case of this situation, where there is a single dominant outbreak across the whole time series, with low numbers of cases in the remaining years. Belém has a more irregular time series of Dengue cases, with multiple outbreaks per year and strong variation in the intensity of outbreaks. In all three cases, the models capture the qualitative changes well, both when selecting the best model for the specific city and when using the best model overall.

**FIG. 2.**
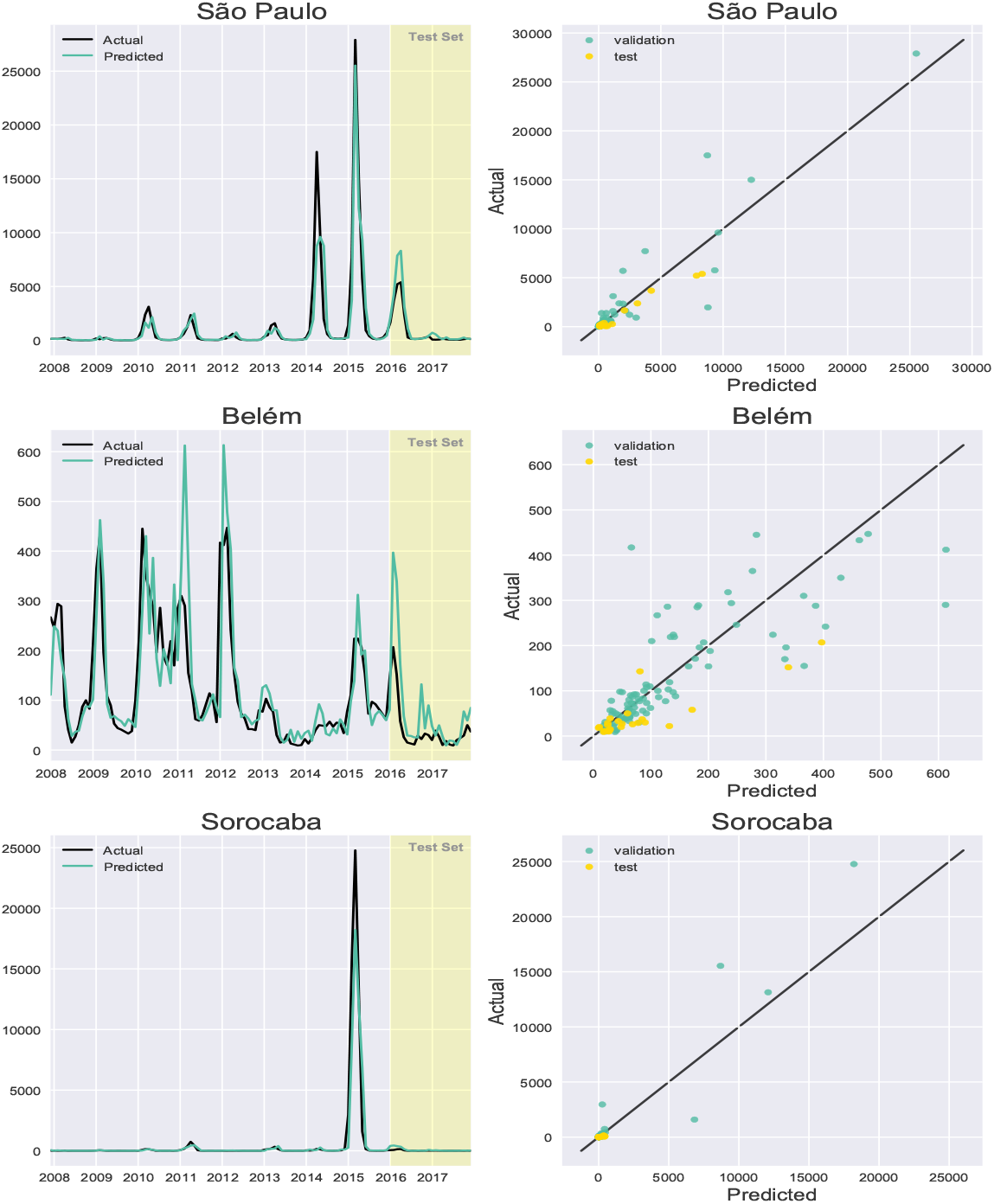
Predictions for sample cities

## IV. DISCUSSION

We compared machine learning algorithms and input feature sets, and show that a Random Forests model trained on eleven lags of historical Dengue cases is effective at forecasting Dengue one month ahead in over 200 Brazilian cities. This model had the lowest aggregate error across cities according to two metrics and was most often chosen as the best-performing model for individual cities.

In some cities, errors on unseen data can be reduced even further by adopting a city-specific approach and selecting one of the other machine learning models. Our study therefore finds that there is no universal model that has the lowest errors in all cities, confirming recent findings in other geographic contexts [47]. However, the increase in the aggregated errors is relatively low when choosing a single model across all cities, and may be justified when considering the increased computational cost of estimating separate models. The most appropriate approach will depend on the intended use case, such as allocating public health resources across cities or developing an early warning system for outbreaks in a specific location.

This study demonstrates the potential of machine learning models to forecast Dengue Fever in Brazil. Our model outperforms existing comparable approaches by achieving a lower forecast error for São Paulo than [31]. Given our data and context, decision tree approaches perform better than neural networks. This finding contributes to better understanding of the role of neural networks in Dengue forecasting, which have not been studied as extensively as other machine learning algorithms [48].

Some limitations of this study must also be considered. We use officially reported statistics of suspected cases, which may include those that are later confirmed to be erroneous. For example, it has been shown that Dengue can be confused with other diseases, such as Zika virus. A study that tested 77 biological samples of a suspected Dengue outbreak near Pernambuco in Northeastern Brazil found that over 40 percent of the patients had actually contracted Zika virus [49]. Therefore, the data and resulting analysis must be viewed in light of these limitations.

A drawback of the machine learning models used in this study is the lack of explainability of the drivers of disease spreading. Effective disease prevention requires an understanding of the causes in addition to the expected number of cases. We can assess the importance of different features for Dengue prediction (e.g. using random forests feature importance or Shapley values), but this would not tell us anything about interventions- which kinds of policies could effectively reduce the disease burden. Detailed exploration of the local drivers in each city can help inform public health policies aimed at suppressing these drivers.

A benefit of using causal discovery for feature selection is the potential for better understanding of the causes of Dengue outbreaks. PCMCI and other causal discovery methods, however, rely on strong assumptions that may limit this benefit in the context of our study. Specifically, PCMCI requires (i) causal sufficiency, (ii) the Causal Markov Condition, and (iii) the Faithfulness assumption [37]. Inferences from the PCMCI analysis must be made carefully when the fulfilment of these assumptions is not proven. For example, the number of both true positives as well as false positives may increase when the stationarity assumption is violated or in the case of long time series [50]. In this study, we use seasonal data and did not include all variables shown to be linked to Dengue Fever, such as human mobility [14] or land use [15]. This does not affect the PCMCI algorithm’s ability to improve predictions, but limits potential causal interpretations. Future work may expand the types of input variables and causal discovery algorithms used to get a better understanding of potential causes of Dengue in Brazilian cities.

In summary, our results show that although Dengue forecasting is of paramount importance, the prediction of the number of cases is not simple and the impact of climate variables depends on the city. A range of factors beyond climate may affect disease spreading. Indeed, since Brazil is among the most unequal countries in the world, it is expected that social indices play a fundamental role in the prediction. Thus, future work may investigate whether social and economic variables can improve dengue forecasting and show which ones are the most important for prediction. In this way, it will be possible to propose methods for epidemic predictions based on the reduction of poverty, carbon emissions, and deforestation.

## ACKNOWLEDGMENTS

This research was supported by grant number 2019/26595-7, São Paulo Research Foundation (FAPESP).

## Notes

### Competing Interest Statement

The authors have declared no competing interest.

